# Molecular Insights into AGS3’s Role in Spindle Orientation: A Biochemical Perspective

**DOI:** 10.1101/2024.07.02.601660

**Authors:** Shi Yu, Jie Ji, Zhijun Liu, Wenning Wang

## Abstract

The intrinsic regulation of spindle orientation during asymmetric cell division (ACD) depends on the evolutionarily conserved protein complex LGN (Pins)/NuMA (Mud)/Gα·GDP. While the role of LGN and its *Drosophila* orthologue Pins is well-established, the function of AGS3, the paralogue of LGN, in spindle orientation during cell division remains controversial. This study substantiates the contentious nature of AGS3’s function through systematic biochemical characterizations. The results confirm the high conservation of AGS3 in its functional structural domains, similar to LGN, and its comparable ability to bind partners including NuMA, Insc and Gα_i3_·GDP. However, in contrast to LGN, AGS3 and the microtubule-binding protein NuMA are unable to form stable hetero-hexamers or higher-order oligomeric complexes that are pivotal for effective regulation of spindle orientation. It was found that this notable difference between AGS3 and LGN stems from the N-terminal sequence preceding the conserved TPR motifs, which spans approximately 20 residues. Furthermore, our findings substantiate the disruptive effect of Insc on the oligomeric AGS3/NuMA complex, while showing no impact on the oligomeric LGN/NuMA complex. Consequently, Insc emerges as an additional regulatory factor that distinguishes the functional roles of AGS3 and LGN, leading to the impairment of AGS3’s ability to actively reorient the mitotic spindle. These results elucidate the molecular basis underlying the observed functional disparity in spindle orientation between LGN and AGS3, providing valuable insights into the regulation of cell division at the molecular level.

## Introduction

In multicellular organisms, asymmetric cell division (ACD) is a fundamental process essential for the generation of diverse cell types (Morrison and Kimble, 2006; D. Konno et al., 2008a). During ACD, the correct alignment of the mitotic spindle plays a critical role in ensuring the unequal segregation of cell fate determinants and production of daughter cells of varying sizes. This process is tightly regulated by a variety of factors, including intrinsic signals and extrinsic cues, as well as mechanical forces (Wu et al., 2008; Lechler and Mapelli, 2021). In the intrinsic signaling regulation of mitotic spindle orientation across various cell types, the protein LGN (Pins in *Drosophila*) plays a pivotal role (Du and Macara, 2004). LGN connects with the apical polarity complex Par3/Par6/aPKC through interactions with Inscuteable (Insc) (Izaki et al., 2006; Kamakura et al., 2013; Williams et al., 2014; Yan et al., 2022), and forms conserved complex LGN/NuMA/Gα_i_. In this complex, the microtubule-binding protein NuMA interacts with motor protein dynein to exert force on astral microtubules, thereby reorienting the spindle (Kiyomitsu and Cheeseman, 2013; Zheng et al., 2013).

Unlike Pins in *Drosophila*, vertebrates possess the paralogs LGN and AGS3, both of which belong to the type II class of activator of G-protein signaling (AGS) family, functioning in G protein signaling activation independently of receptors (Bernard et al., 2001; Blumer et al., 2012). AGS3 shares high sequence similarity with LGN, characterized by an N-terminal TPR (tetratricopeptide repeats) domain containing eight TPR motifs and a C-terminal GoLoco domain containing four GoLoco (GL) motifs (Fig. 1A) (Groves et al., 2007; Jia et al., 2012). In addition to its involvement in GPCR signaling regulation, AGS3 has been linked to various cellular functions, including neuronal differentiation, autophagy, protein transport, and brain adaptation mechanisms associated with the renal injury response (Pattingre et al., 2003; Groves et al., 2007; Blumer et al., 2008; Bowers et al., 2008; Fan et al., 2009; Regner et al., 2011).

**Figure 1.**
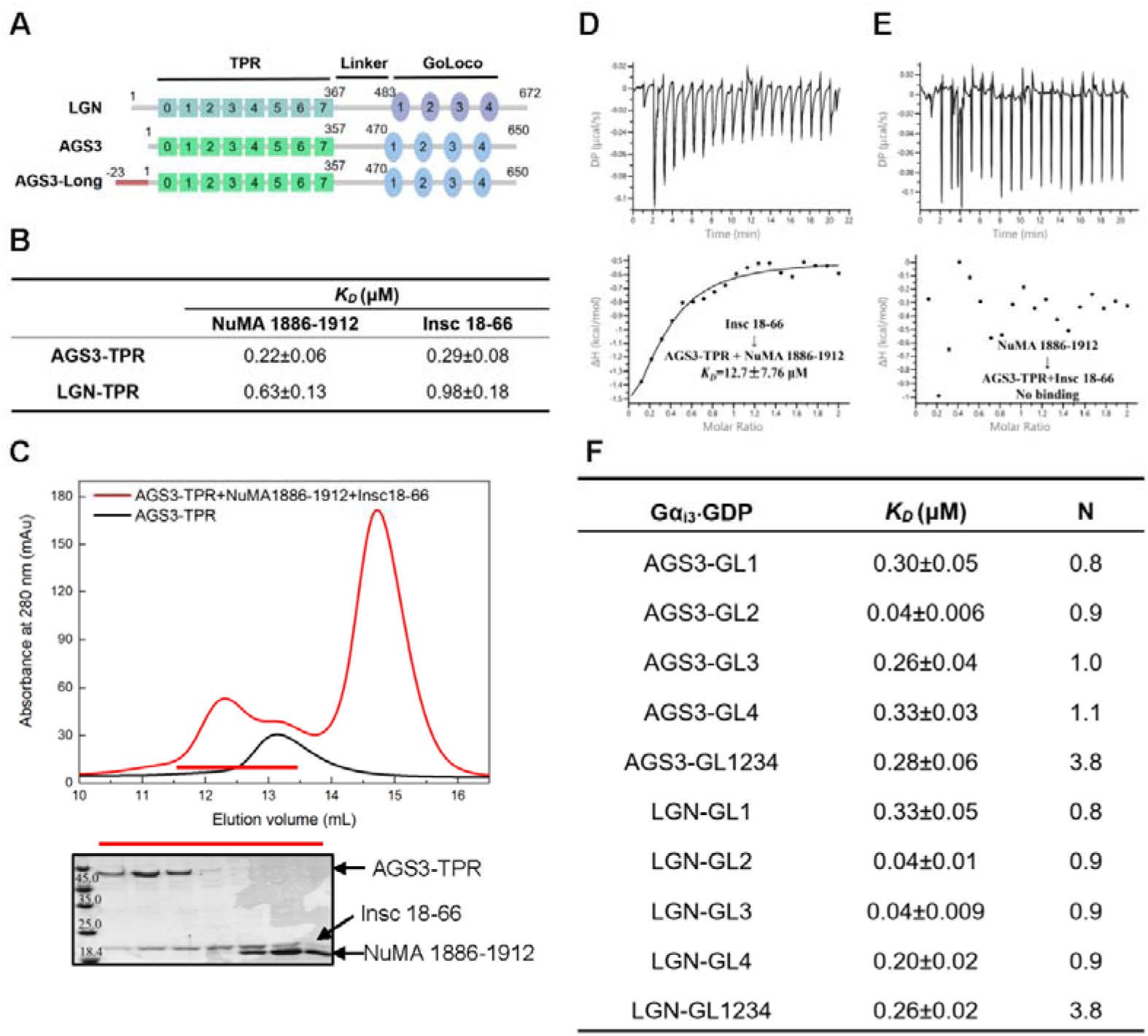
AGS3-TPR and AGS3-GL interact with their partners similarly to LGN. (A) The domain organizations of LGN and AGS3 isoforms. (B) The ITC-based binding affinities of the TPR domains of AGS3/LGN to Insc and NuMA. The raw data and fits are shown in the Supplementary material Figure S1. (C) The SEC analysis of the mixture of AGS3-TPR, NuMA (1886-1912) and Insc (18-66) at a 1:2:2 ratio. (D) ITC measurement of the binding of Insc (18-66) to the mixture of AGS3-TPR and NuMA (1886-1912). (E) ITC measurement of the binding of NuMA (1886-1912) to the mixture of AGS3-TPR and Insc (18-66). (F) ITC-based measurements of the binding affinities of Gα_i3_·GDP to GL motifs and GoLoco domains of AGS3/LGN. The raw data and fits are shown in the Supplementary material Figure S2.

Nonetheless, the role of AGS3 in spindle orientation remains controversial. Early research suggested that AGS3 perturbs planar spindle orientation in apical divisions of neuronal progenitor cells (Sanada and Tsai, 2005). However, another study observing RNAi targeting AGS3 did not reveal a more planar spindle orientation phenotype (Daijiro Konno et al., 2008b). Subsequently, Saadaoui et al. discovered that despite its conserved interactions with NuMA and Gα_i_ *in vitro*, AGS3 cannot substitute LGN for spindle orientation function *in vivo* (Saadaoui et al., 2017). More recently, Descovich et al. found that AGS3 promotes planar divisions by counteracting LGN’s ability in promoting and maintaining vertical divisions, implying opposing roles for these two vertebrate paralogs in regulating epidermal spindle orientation and differentiation during development (Descovich et al., 2023). These observations underscore the need to understand the structural foundation underlying the distinct functional roles of LGN and AGS3 in spindle orientation. While there have been some studies investigating AGS3’s domain interactions (Bernard et al., 2001; Adhikari and Sprang, 2003; Sanada and Tsai, 2005; Saadaoui et al., 2017; Wavreil and Yajima, 2020; Yip et al., 2020), a comprehensive comparison of LGN and AGS3 in terms of their domain structures and protein-protein interactions, especially with quantitative evaluation, remains lacking.

In this study, we conducted a comprehensive biochemical characterization to parallelly investigate the inter- and intramolecular interactions of AGS3 and LGN. Our findings indicate that the binding affinities of the TPR domain to NuMA and Insc, as well as the GoLoco domain with Gα_i3_·GDP, are very similar between AGS3 and LGN. Additionally, we compared the intramolecular interactions and the opening of the auto-inhibited conformation of AGS3 and LGN, revealing analogous behaviors in both proteins. Furthermore, we evaluated the capabilities of AGS3 and LGN to form hetero-oligomer complexes with NuMA, revealing that the multimeric AGS3/NuMA complex exhibits reduced stability compared to LGN/NuMA complex. Moreover, the AGS3/NuMA complex is susceptible to disruption by Insc, highlighting a potential regulatory mechanism that differs from LGN/NuMA interactions. These findings provide valuable insights into the molecular mechanisms underlying the functional disparities observed in spindle orientation between AGS3 and LGN. By elucidating these differences at the biochemical level, our study contributes to a better understanding of how these paralogs exert distinct roles in regulating cellular processes, particularly in the context of spindle orientation and asymmetric cell division.

## Results

### The TPR and GoLoco domains of AGS3 display binding patterns similar to those of LGN with their respective partners

Previous studies have demonstrated that the TPR domain of AGS3 can interact with NuMA and Insc similar to LGN (Izaki et al., 2006; Yuzawa et al., 2011; Saadaoui et al., 2017). However, there is a lack of systematic quantification of these interactions. The interactions of AGS3-TPR (1-405) with NuMA and Insc were initially characterized using isothermal titration calorimetry (ITC). The *K_D_*values for the binding of AGS3-TPR with Insc (18-66) and NuMA (1886-1912) were determined to be 0.29 μM and 0.22 μM, respectively, analogous to those of LGN-TPR (1-409) (Fig. 1B and Fig. S1A-S1D). Therefore, the TPR domain of AGS3 exhibits comparable affinities for NuMA and Insc as LGN. The mutual exclusivity of NuMA and Insc binding to LGN-TPR has been well-documented, with Insc having the capability to supplant NuMA in the binding process. (Culurgioni et al., 2011; Zhu et al., 2011). Here, we examined whether AGS3-TPR also exhibits a preference for binding to Insc over NuMA. To address this, we mixed AGS3-TPR with Insc (18-66) and NuMA (1886-1912) at a 1:2:2 molar ratio, and characterized the mixture using size exclusion chromatography (SEC) and SDS-PAGE. The results showed that only Insc (18-66) was present in the complex with AGS3-TPR (Fig. 1C), suggesting a preferentially binding of AGS3-TPR to Insc over NuMA. A similar analysis of the LGN-TPR/Insc/NuMA mixture yielded consistent results with the previous study, showing that LGN preferentially binds to Insc (Fig. S1E). Additionally, ITC measurements demonstrated that Insc could bind the 1:1 mixture of AGS3-TPR and NuMA, whereas NuMA could not bind the 1:1 mixture of AGS3-TPR and Insc (Fig. 1D and 1E). Taken together, these results indicate that the TPR domain of AGS3 shares similar characteristics of partner interactions with LGN-TPR.

Similar to LGN-GoLoco, the GoLoco domain of AGS3 has been demonstrated to interact with Gα_i3_·GDP or Gα_i1_·GDP (Bernard et al., 2001; Adhikari and Sprang, 2003; Sanada and Tsai, 2005; Yip et al., 2020). Detailed biochemical analyses of the interactions between AGS3-GoLoco and Gα_i1_·GDP have revealed binding affinities that resemble those observed between LGN-GoLoco and Gα_i_·GDP (Adhikari and Sprang, 2003; Jia et al., 2012). In this study, we specifically investigated the binding affinities of AGS3-Goloco to Gα_i3_·GDP. The results showed that each GL motif (~ 34 residues) of AGS3 exhibited robust binding to Gα_i3_·GDP, displaying binding affinities comparable to those observed for LGN’s GL motifs (Fig. 1F and Fig. S2A-H). Furthermore, the full-length GoLoco domains (GL1234) of AGS3 and LGN also showed similar binding affinities to Gα_i3_·GDP, consistent with a 1:4 binding stoichiometry (Fig. 1F and Fig. S2I-J). Therefore, both AGS3- and LGN-GoLoco domains can bind four molecules of Gα_i3_·GDP simultaneously, with apparent binding affinities resembling to those observed for individual GL motifs (Fig. 1F).

### AGS3 forms auto-inhibited conformation through intramolecular interactions

LGN adopts an auto-inhibited conformation through intramolecular interaction between its TPR and GoLoco domains(Pan et al., 2013). In this study, we characterized the intramolecular interactions within AGS3. Result from ITC measurement revealed a binding affinity of 0.17 μM between AGS3-TPR and AGS3-GoLoco (Fig. 2A). To examine the auto-inhibition, fusion proteins of AGS3 and LGN were constructed by replacing the linker sequences between the TPR and GoLoco domains with eight pairs of GS amino acids. The resultant fusion proteins were designated as FL-AGS3 and FL-LGN (Fig. 2B). Subsequent ITC experiments demonstrated that both FL-AGS3 and FL-LGN lost the ability to bind their isolated GoLoco domains (Fig. 2C and 2D), indicating that they adopt an autoinhibited conformation characterized by strong intramolecular interactions between the TPR and GoLoco domains.

**Figure 2.**
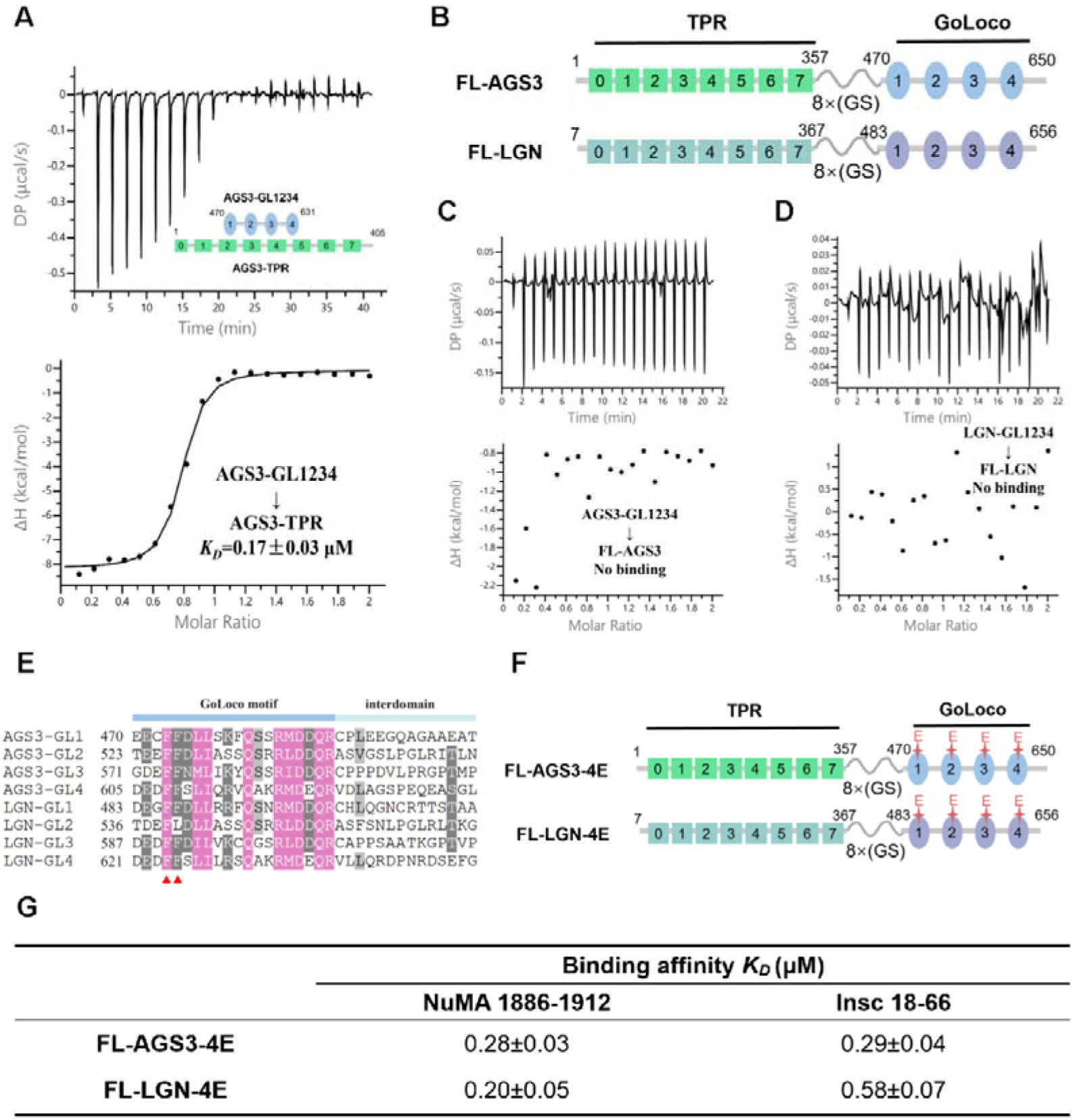
Intramolecular interactions and autoinhibited conformation of AGS3. (A) The ITC measurement of the interaction between AGS3-TPR and AGS3-GL1234. (B) Schematic diagram of the domain organization of FL-AGS3 and FL-LGN. (C) The ITC measurement of the interaction between AGS3-GL1234 to FL-AGS3. (D) The ITC measurement of the interaction between LGN-GL1234 to FL-LGN. (E) Sequence alignment of the GL motifs of AGS3 and LGN. The “interdomain” regions are the variable sequences that separate the conserved part of each GL motifs. The red triangles indicate the conserved phenylalanines. (F) Schematic diagram of the domain organization of FL-AGS3-4E and FL-LGN-4E. (G) ITC-based binding affinities of FL-AGS3-4E and FL-LGN-4E mutants to Insc (18-66) and NuMA (1886-1912). The raw data and fits are shown in the Supplementary material Figure S3.

The sequence analysis of the GL motifs reveals a high level of conservation of key hydrophobic amino acids (F486, F487, F539, F591, and F624) essential for TPR interactions in both AGS3 and LGN (Fig. 2E) (Pan et al., 2013). Based on this observation, we hypothesized that substitution of these Phe with charged amino acids in all four GL motifs could disrupt the auto-inhibition. Mutant proteins, where all four Phe residues were replaced by Glu, were designated as FL-AGS3-4E and FL-LGN-4E (Fig. 2F). Subsequent ITC results indicated that both FL-AGS3-4E and FL-LGN-4E mutant proteins exhibited strong binding affinities with Insc or NuMA, similar to the affinities observed for the isolated TPR domain (Fig. 2G and Fig. S3). These data suggest that, akin to LGN, the 4E mutation disrupts intramolecular interactions in AGS3, thereby releasing the autoinhibited conformation.

### Both Insc and NuMA could open the auto-inhibited conformation of AGS3

To elucidate the influence of Insc and NuMA on the auto-inhibited conformation, we measured the binding affinities between Insc/NuMA and the FL-AGS3/LGN proteins employing ITC. The results showed that Insc (18-66) or NuMA (1808-2001) could directly bind to FL-AGS3/FL-LGN (Table 1 and Fig. S4). Moreover, the binding affinities exhibited no substantial augmentation in the presence of Gα_i3_·GDP (Table 1 and Fig. S4). This suggests that both Insc and NuMA have the ability to perturb the auto-inhibited conformation of AGS3, and Gα_i3_·GDP does not play a requisite role in facilitating Insc/NuMA’s interaction with AGS3.

**Table 1.**
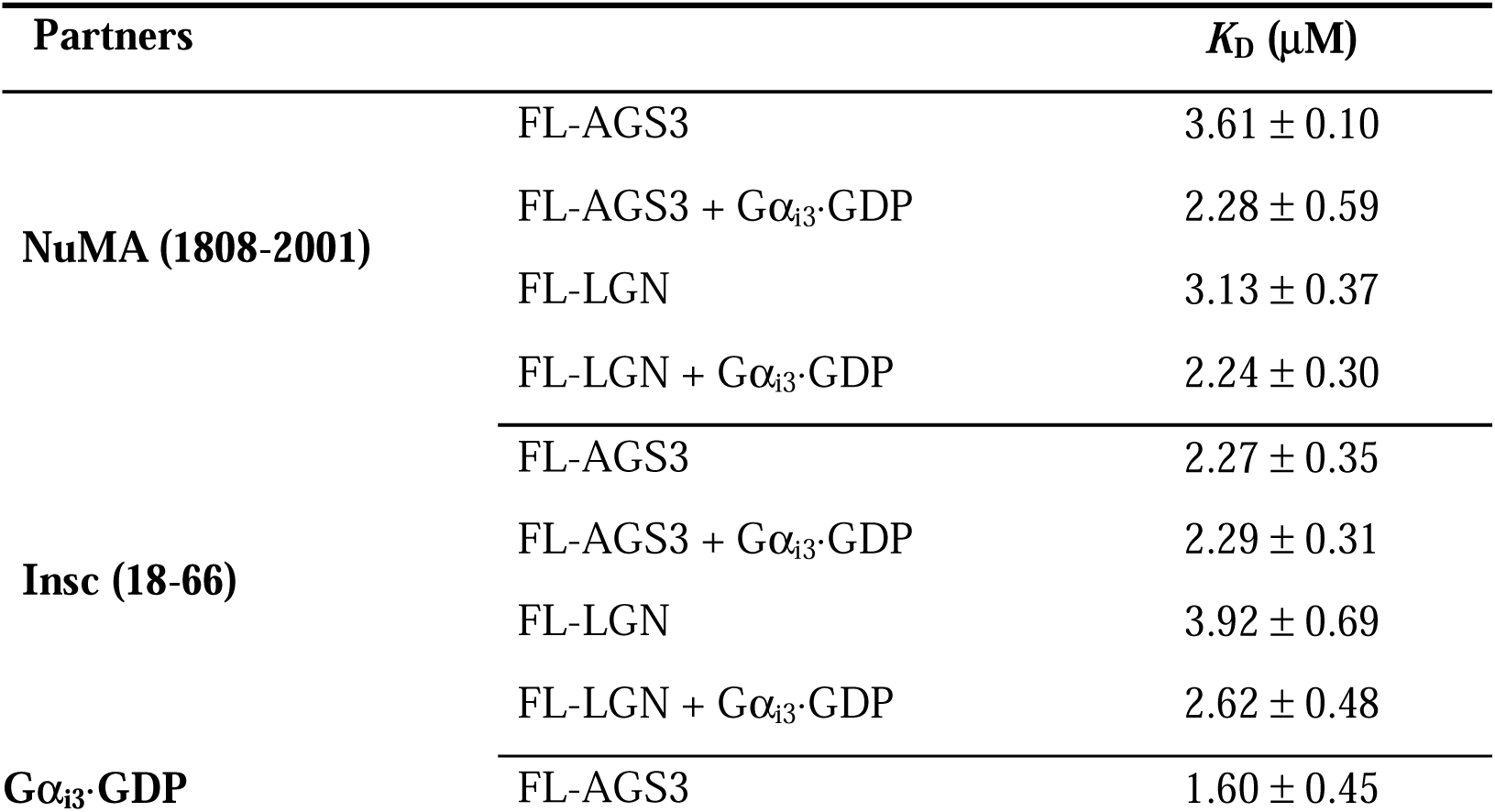
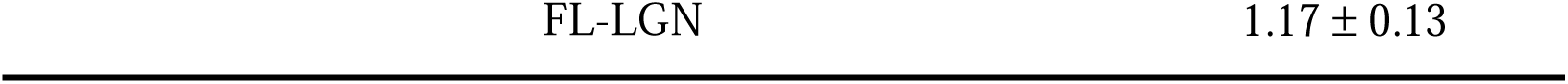
Binding affinities of FL-AGS3/LGN to their partners.

We additionally characterized the interaction between Gα_i3_·GDP and FL-AGS3/FL-LGN. The results indicate that Gα_i3_·GDP binds to FL-AGS3 and FL-LGN with comparable affinities (Table 1, Fig. S5). These suggest that Gα_i3_·GDP is capable of binding to the auto-inhibited states of AGS3 and LGN without the assistance of other proteins.

### Long-form AGS3-TPR with additional N-terminal residues could form oligomeric complex with NuMA

Recent structural studies have revealed that LGN and NuMA form specific hetero-hexamers and also bind to dimeric full-length NuMA, thus generating an extended subcortical protein network (Pirovano et al., 2019). These oligomeric LGN/NuMA complexes facilitate microtubule capture at the cell cortex, essential for spindle orientation in epithelial cells (Pirovano et al., 2019). The ability of the TPR domain of LGN to form higher-order oligomeric complexes with NuMA appears to be critical for its spindle orientation function. However, it remains unclear whether AGS3 and NuMA can also form analogous oligomeric complex. According to the literature findings, the shortest segments capable of forming hetero-hexamers are LGN (7-367) and NuMA (1847-1914) fragments (Pirovano et al., 2019). Consistent with these reports, our SEC results indicated that LGN-TPR (1-409)/NuMA (1847-1914) formed a complex (the peak at ~10.5 mL) with a molecular weight (MW) well above 150 kDa, lager than that of the binary LGN (15-350)/NuMA (1847-1914) complex (the peak at ~12.4 mL) (Fig. 3A and 3B). Static light scattering (SLS) experiment further confirmed that the MW of the LGN-TPR/NuMA (1847-1914) complex was 227 kDa (Fig. 3C), consistent with a hetero-hexamer formation with a 3:3 binding ratio. What’s more, the shorter NuMA fragment (NuMA 1886-1912) in conjunction with LGN-TPR failed to form an oligomer with MW higher than 1:1 complex (Fig. S6A). These results highlight the necessity of the N-terminal 14 residues of LGN and the extended NuMA fragment (1847-1914) for the formation of LGN/NuMA hetero-hexamer.

**Figure 3.**
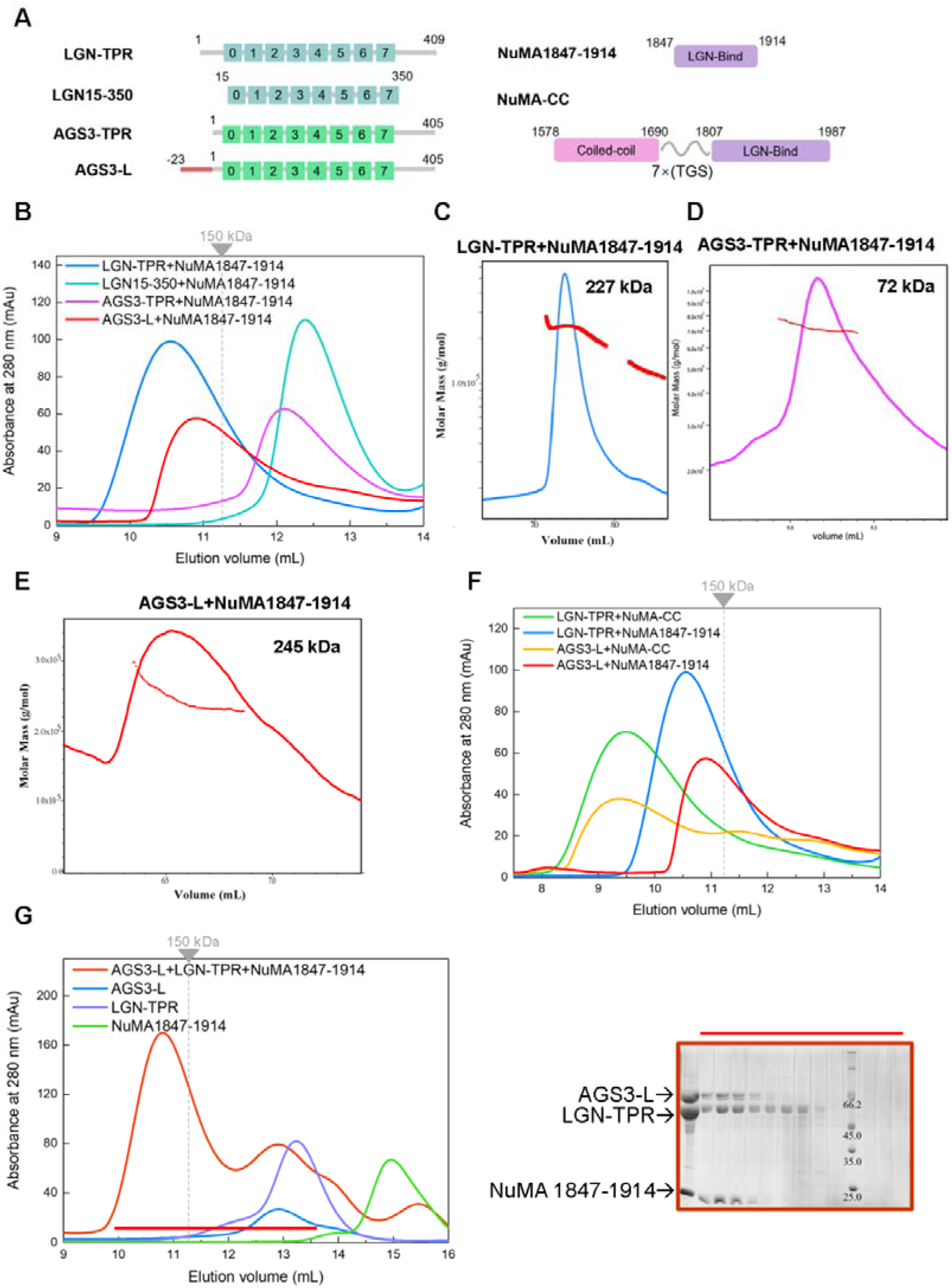
Long-form AGS3-TPR with additional N-terminal residues could form multimeric complex with NuMA. (A) Schematic diagram of the domain organization of various fragments of AGS3, LGN and NuMA. (B) SEC analyses of the complexes formed between LGN-TPR, AGS3-TPR or AGS3-L with NuMA (1847-1914). The dashed line indicates the elution volume of a 150 kDa globular protein marker. Static light scattering characterization of the molecular weight of the LGN-TPR/NuMA (1847-1914) complex (C), AGS3-TPR/NuMA (1847-1914) complex (D) and AGS3-L/NuMA (1847-1914) complex (E). (F) SEC analyses of the complexes formed between LGN-TPR or AGS3-L with NuMA-CC. The dashed line indicates the elution volume of a 150 kDa globular protein marker. (G) SEC analyses of the complex formed between AGS3-L, LGN-TPR and NuMA (1847-1914). Coomassie-stained SDS-PAGE to the right shows the protein composition of corresponding elution profiles.

In contrast, the SEC analysis of the AGS3-TPR/NuMA (1847-1914) mixture (the peak at ~12.1 mL) exhibited a MW below 150 kDa, indicating the absence of an oligomeric complex (Fig. 3B). Further SLS experiment confirmed the MW of AGS3-TPR/NuMA (1847-1914) as 72 kDa, consistent with 1:1 binding ratio (Fig. 3D). Sequence alignment of AGS3 and LGN reveals that AGS3-TPR (1-405) corresponds to LGN (13-409) (Fig. S6C), which lacks the critical N-terminal residues necessary for LGN hexamer formation. We hypothesized that a longer isoform with additional N-terminal residues might be required for AGS3 to form a multimeric complex with NuMA. Therefore, we purified a longer form of AGS3-TPR containing additional 23 residues at the N-terminus (Uniprot entry **<u>Q86YR5-1</u>**), denoted as AGS3-L (-23-405, with the N-terminal residues labeled from −23 to −1 to align with the sequence numbering of the short isoform) (Fig. 3A), and then examined its mixture with NuMA (1847-1914).

AGS3-L exhibited similar binding affinities to NuMA and Insc compared to AGS3-TPR (Fig. S7). SEC analysis revealed that, similar to LGN-TPR, AGS3-L formed a complex with NuMA (1847-1914) (the peak at ~10.9 mL) with a MW significantly exceeding 150 kDa (Fig. 3B). SLS measurement further verified that the MW of the complex as 245 kDa (Fig. 3E), indicative of a 3:3 hetero-hexamer complex. Additionally, when the NuMA fragment was truncated to (1886-1912), AGS3-L was no longer able to form a hexamer with NuMA (Fig. S6B). These findings underscore the requirement for the additional N-terminal residues in AGS3-L for efficient formation of a hetero-hexamer with NuMA.

To gain insights into the architecture of the AGS3-L/NuMA oligomeric complex, we built a structural model of the AGS3-L/NuMA (1847-1914) hexametric complex using homology modeling (Webb and Sali, 2016) based on the crystal structure of LGN/NuMA hexamer (PDB: 6HC2). The model reveals two critical structural characteristics necessary for hexamer formation (Fig. S8A). First, the additional 23 residues at the N-terminus of AGS3-L fold into a helix (colored in slate blue in Fig. S8B), and forms four-helix bundle with the TPR8 motif from the adjacent AGS3-L (Fig. S8B). Secondly, the N-terminal region of NuMA (1847-1914) interacts with the neighboring AGS3-L and contributes to stabilizing the hexamer structure (Fig. S8C). Overall, the structural model of AGS3-L/NuMA hexamer verifies the necessity of the N-terminal extension of the long-form AGS3 in forming oligomeric complex with NuMA.

Full-length NuMA is known to form dimers through its coiled-coil domain, and previous study have demonstrated that dimeric NuMA facilitates the formation of higher-than-hexamer LGN/NuMA complex (Pirovano et al., 2019). To explore this further, we constructed a fusion protein connecting NuMA’s coiled-coil domain (1578-1690) with its LGN-binding domain (1807-1987) via a 7(TGS) linker, designated as NuMA-CC (i.e. 1578-1690-7(TGS)-1807-1987) (Fig. 3A). In agreement with the previous study (Pirovano et al., 2019), SEC analysis showed that the size of the LGN-TPR/NuMA-CC complex (the peak at ~9.5 mL) was significantly larger than that of LGN-TPR/NuMA (1847-1914) (Fig. 3F), suggesting the formation of a higher-order multimeric complex beyond a hexamer. Similarly, SEC analysis indicated that the MW of the complex formed by AGS3-L and NuMA-CC (the peak at ~9.3 mL) was also much larger than that of AGS3-L/NuMA (1847-1914) (Figure 3F), indicating that AGS3-L can form a high-order hetero-oligomer with NuMA-CC as well. These results underscore the capacity of both LGN and AGS3 to engage in higher-order complex formation with NuMA.

### Competitive binding between AGS3 and LGN for NuMA

Subsequently, we investigated the competitive binding between AGS3 and LGN for NuMA. A mixture containing AGS3-L, LGN-TPR and NuMA (1847-1914) in a 1:1:1 ratio was analyzed using SEC to assess the composition of the complex formed (the peak at ~10.8 mL) (Fig. 3G). The results confirmed the presence of all three components, indicating that both LGN and AGS3 can engage with NuMA (Fig. 3G). Furthermore, a GST pull down experiment was conducted to evaluate the binding of AGS3-L and LGN-TPR to NuMA (1808-2001) at various concentration ratios. The results demonstrated that AGS3-L and LGN-TPR displayed comparable binding abilities for NuMA (Fig. S9). Moreover, an excess of AGS3-L over LGN-TPR resulted in the displacement of LGN-TPR and complete occupancy of NuMA (Fig. S9A). Conversely, an excessive amount of LGN-TPR predominantly led to the formation of LGN-TPR/NuMA complex (Fig. S9B). These findings highlight the competitive nature of AGS3 and LGN for binding to NuMA, influenced by their relative concentrations.

### Full-length AGS3/LGN has less capacity to form oligomeric complex with NuMA

Both LGN and AGS3 exhibit strong intramolecular interactions in their full-length forms, which may impact the assembly of high-order oligomers with NuMA. In order to investigate the binding of full-length LGN and AGS3 to NuMA, we utilized the FL-LGN fusion proteins mentioned above and engineered a long-form AGS3 protein, which includes the N-terminal 23 residues (i.e., -23-357-8(GS)-470-650), designated as FL-AGS3-L (Fig. 4A). Additionally, the preceding data indicated that NuMA (1808-2001) can bind to FL-LGN and FL-AGS3 in the absence of Gα_i3_·GDP. Therefore, we conducted SEC analysis to examine the mixture of FL-LGN or FL-AGS3-L with NuMA (1808-2001). The results showed that FL-LGN forms a 1:1 complex with NuMA (1808-2001) (the peak at ~11.5 mL) (Fig. 4B). This contrasts with LGN-TPR, which forms 3:3 hetero-hexamer with NuMA (Fig. 3B), suggesting that intramolecular interactions within FL-LGN hinder the formation of multimeric complex. Introducing the 4E mutation in FL-LGN to disrupt these intramolecular interactions, the resultant FL-LGN-4E protein leads to the formation of a higher MW complex (the peak at ~10.3 mL) when mixed with NuMA (1808-2001) (Figure 4a). The SLS experiment revealed that the MW of the FL-LGN-4E/NuMA (1808-2001) complex was 196 kDa, indicative of a 2:2 hetero-tetramer (Fig. 4C). These results suggest that only in the open conformational state can the FL-LGN form a multimeric complex with NuMA, and the GoLoco domain impedes the formation of high-order oligomer even without intramolecular interactions.

**Figure 4.**
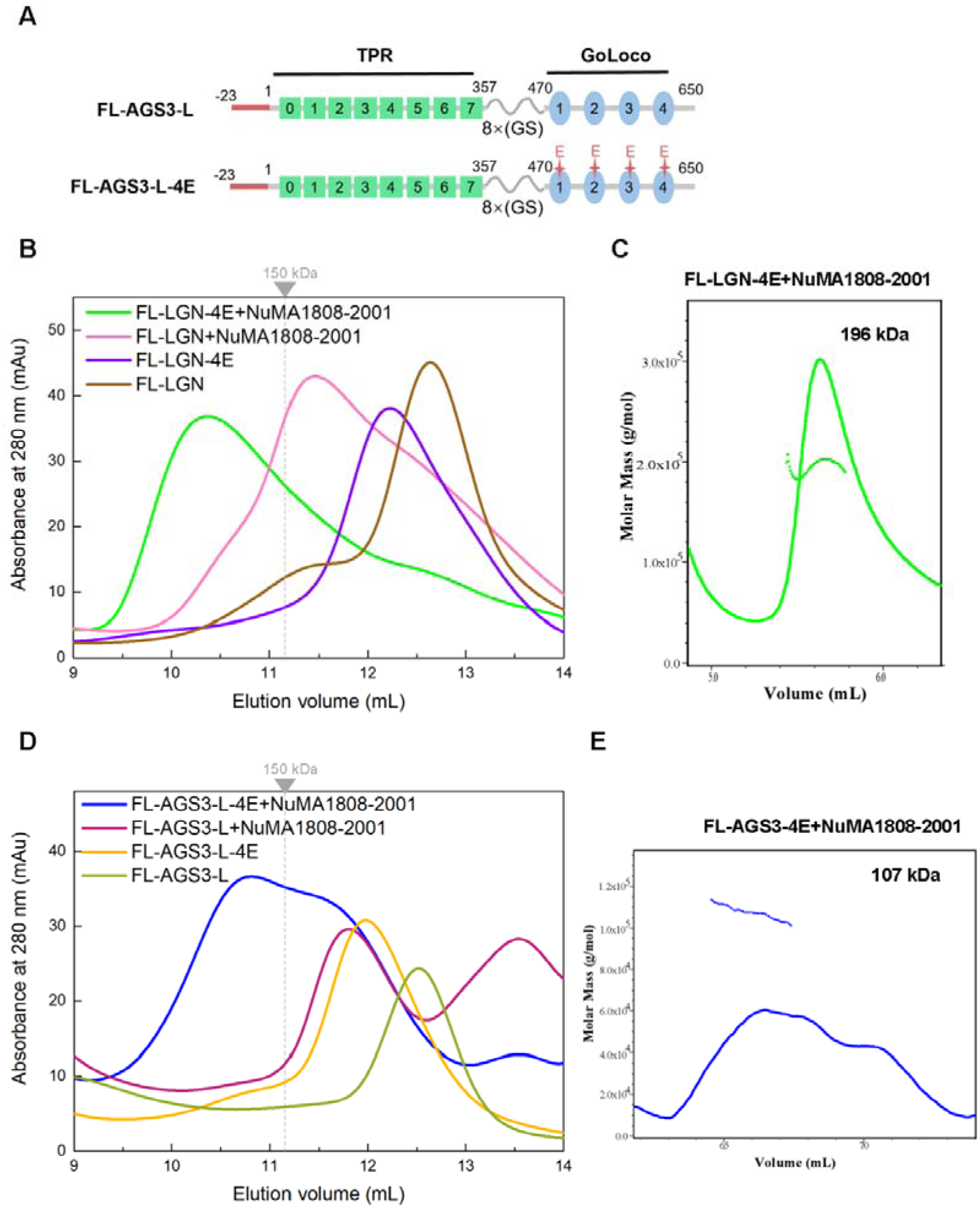
Full length LGN and AGS3 have lower capacity to form oligomeric complex with NuMA. (A) Schematic diagram of the domain organization of long isoform of FL-AGS3-L and FL-AGS3-L-4E. (B) SEC analyses of the complexes formed between FL-LGN or FL-LGN-4E with NuMA (1808-2001). The dashed line indicates the elution volume of a 150 kDa globular protein marker. (C) SLS characterization of the molecular weight of the FL-LGN-4E/NuMA (1808-2001) complex. (D) SEC analyses of the complexes formed between FL-AGS3-L or FL-AGS3-L-4E with NuMA (1808-2001). (E) SLS characterization of the molecular weight of the FL-AGS3-L-4E/NuMA (1808-2001) complex.

For AGS3, the SEC profile demonstrated that the FL-AGS3-L fusion protein forms a 1:1 complex with NuMA (1808-2001) (the peak at ~11.8 mL) (Fig. 4D). Upon introducing the FL-AGS3-L-4E mutant to disrupt the intramolecular interaction, the elution volume of complex shifted to a smaller volume (the peak at ~10.8 mL) (Fig. 4D). However, the peak was broad, and the MW was obviously smaller compared to that of the FL-LGN-4E/NuMA complex (Fig. 4D). Subsequent SLS experiment indicated a MW of 107 kDa, implying primarily a 1:1 ratio complex (Fig. 4E). Consequently, these results indicate that FL-AGS3-L is unable to form a stable multimeric complex with NuMA, even in the open conformation. This suggests that AGS3 has a lower capacity than LGN to engage in high-order complex formation with NuMA.

### The impact of Insc on the multimeric complex formed between LGN/AGS3 and NuMA

We have demonstrated that the TPR domains of both AGS3 and LGN exhibit a preference for binding with Insc over NuMA. The preceding findings suggest that LGN-TPR and AGS3-L can form multimeric complexes with NuMA. These observations prompted us to investigate whether the TPR domain of AGS3 or LGN retains this preference in the context of multimeric complex formation with NuMA. To address this question, LGN-TPR or AGS3-L proteins were mixed with either NuMA (1847-1914) or NuMA-CC along with Insc (18-66) protein in a 1:2:2 ratio, and the resulting mixtures were subsequently analyzed by SEC. The results revealed that the introduction of Insc leads to a slight reduction in the quantity of LGN/NuMA hetero-hexamer complexes (the peak at ~10.8 mL) (Fig. 5A), indicating that Insc partially disrupted the LGN/NuMA hetero-hexamers. This suggests that Insc competes, to some extent, with NuMA (1847-1914) for binding to LGN. Similarly, the high-order LGN-TPR/NuMA-CC complex (the peak at ~9.5 mL) was also partly disrupted by the addition of Insc (18-66) (Fig. 5B), albeit to a lesser degree compared to NuMA (1847-1914) (Fig. 5A).

**Figure 5.**
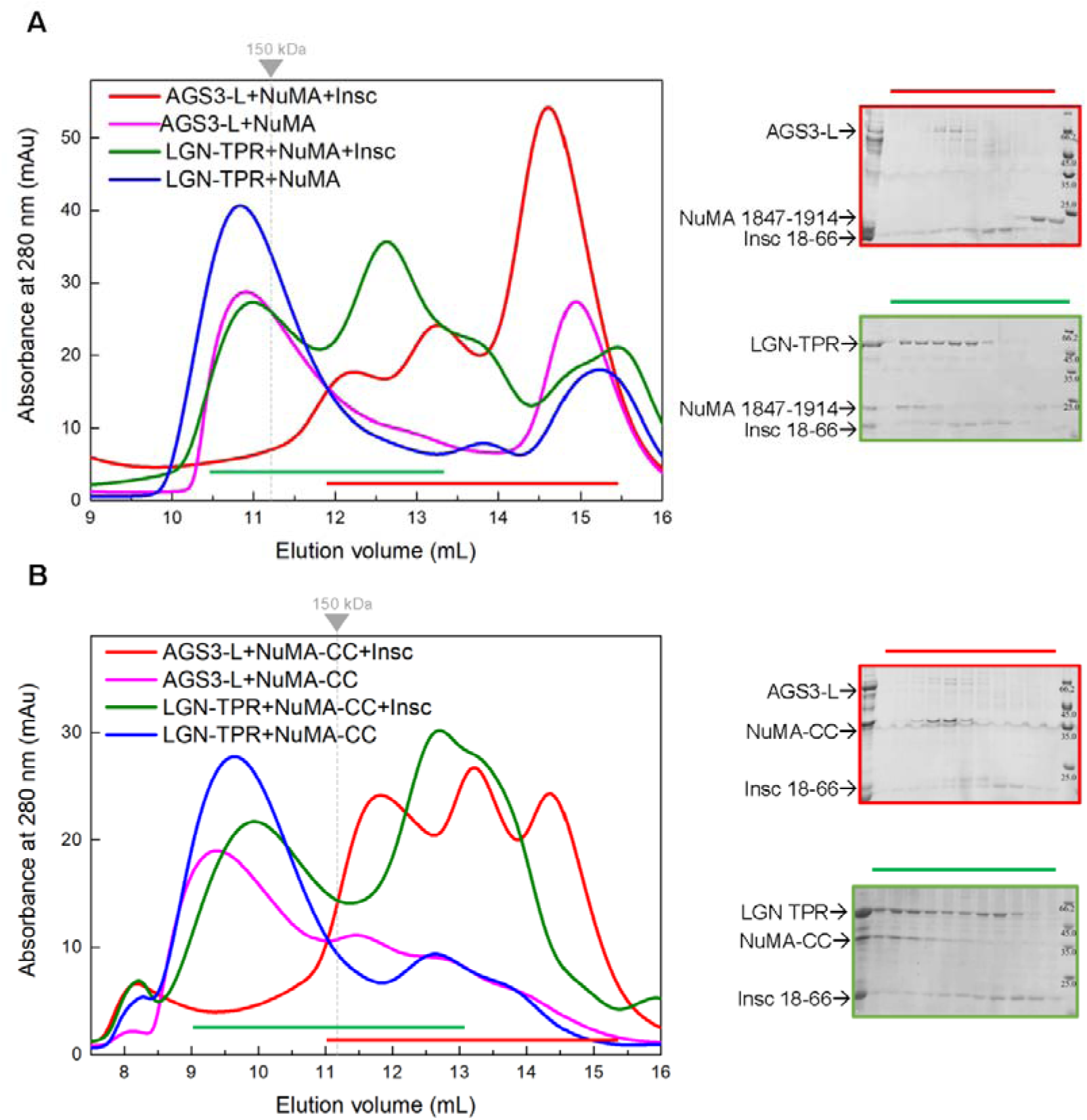
The impact of Insc on the oligomeric complexes formed between LGN or AGS3 with NuMA. (A) SEC analyses of the complex formed between LGN-TPR or AGS3-L with NuMA (1847-1914) in the presence and absence of Insc (18-66). Coomassie-stained SDS-PAGE to the right show the protein composition of two elution profiles. (B) SEC analyses of the complex formed between LGN-TPR or AGS3-L with NuMA-CC in the presence and absence of Insc (18-66). Coomassie-stained SDS-PAGE to the right show the protein composition of two elution profiles.

In contrast, the multimeric complexes AGS3-L/NuMA (1847-1914) (the peak at ~10.9 mL) and AGS3-L/NuMA-CC (the peak at ~9.3 mL) were nearly completely disrupted by Insc (18-66) (Fig. 5A and 5B). A significant portion of AGS3-L instead formed a complex with Insc (18-66) (the peak detected at ~12.1 mL) (Fig. 5A and 5B). These findings suggest that the high-order oligomeric complexes formed between AGS3-L and NuMA are less stable than those between LGN and NuMA. Therefore, these results support the conclusion that Insc maintains a preference for binding to the TPR domain of AGS3, even in the context of oligomeric complex formation with NuMA.

### The N-terminal sequences determine the different capacities of AGS3 and LGN in forming oligomeric complex with NuMA

We hypothesized that the reduced stability of the oligomeric AGS3-L/NuMA complexes might be attributed to the less conserved N-terminal residues in the AGS3-L sequence, which exhibit lower homology with the corresponding region of the LGN (Fig. S6C). To investigate the significance of these N-terminal sequences in LGN and AGS3 for their ability to oligomerize with NuMA, we engineered a chimeric fusion protein. This construct substituted the N-terminal segment (-23~-1) of AGS3-L with residues 1-12 from the LGN, resulting in the construct AGS3^LGN^ (i.e., LGN (1-12)-AGS3 (1-405)) (Fig. 6A). In a reciprocal approach, we also generated an LGN-TPR fusion protein, LGN^AGS3^ (i.e., AGS3 (-23~-1)-LGN (13-409)), where the N-terminal 1-12 amino acids of LGN were replaced with the N-terminal segment (-23~-1) from AGS3 (Fig. 6A).

**Figure 6.**
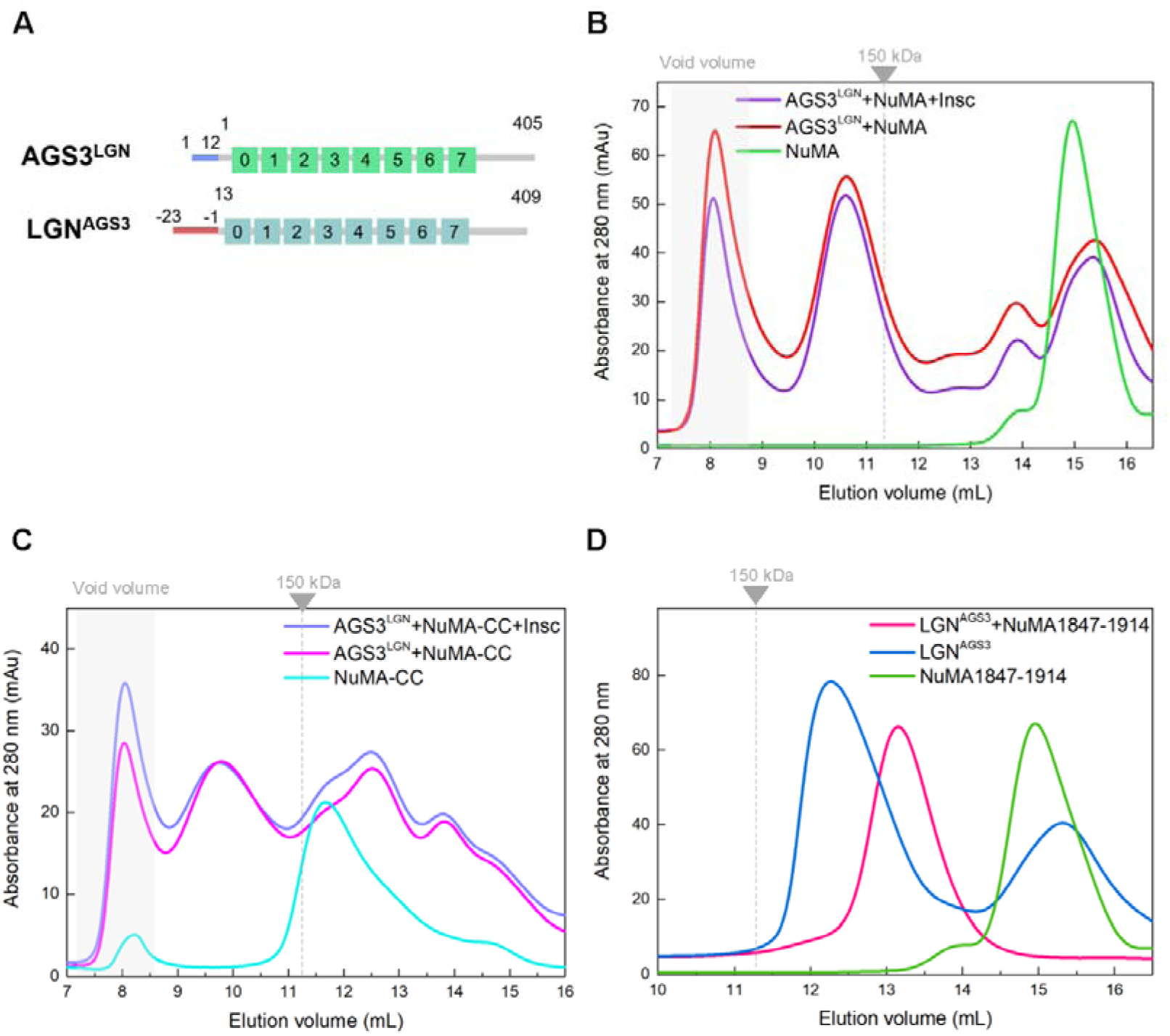
The N-terminal sequences determine the different capacities of AGS3 and LGN in forming multimeric complex with NuMA. (A) Schematic diagram of the domain organization of chimeric proteins AGS3^LGN^ and LGN^AGS3^. (B) SEC analyses of the complex formed between AGS3^LGN^ and NuMA (1847-1914) in the presence and absence of Insc (18-66). The gray box shows the void volume. (C) SEC analyses of the complex formed between AGS3^LGN^ and NuMA-CC in the presence and absence of Insc (18-66). The gray box shows the void volume. (D) SEC analyses of the complex formed between LGN^AGS3^ and NuMA (1847-1914).

Subsequent SEC analyses were conducted to examine the interactions between AGS3^LGN^ or LGN^AGS3^ with NuMA (1847-1914). The results demonstrated that AGS3^LGN^ exhibited the ability to form a stable multimeric complex with NuMA (1847-1914), and the presence of Insc did not disrupt this interaction (Fig. 6B). This behavior markedly contrasts with AGS3-L, which failed to maintain a stable hetero-hexamer complex with NuMA in the presence of Insc (Fig. 5A). Similarly, for the AGS3^LGN^/NuMA-CC complex, the addition of Insc also produced negligible interference effects (Fig. 6C).

Conversely, unlike LGN-TPR, the LGN^AGS3^ protein was incapable of forming a hetero-hexamer complex with NuMA, instead forming a 1:1 complex (Fig. 6D). These experimental findings reinforce the notion that the non-conserved N-terminal residues preceding the TPR motifs in LGN and AGS3 play a critical role in determining their ability to form high-order multimeric complex with NuMA.

## Discussion

In this research, we conducted a systematic examination aimed at discerning the structural and functional disparities between AGS3 and LGN, with a particular emphasis on biochemical analysis of the intra- and inter-molecular interactions of the TPR and GoLoco domains. Our findings revealed that the isolated TPR or GoLoco domains of the both proteins exhibit comparable binding affinities towards their respective partners, which include NuMA, Insc, and Gα_i3_·GDP. Minimal variations were observed in intramolecular interactions and in the release of their autoinhibited conformations. The primary distinction observed between AGS3 and LGN lies in their respective abilities to form oligomeric complexes with NuMA. The short-form AGS3-TPR without the N-terminal 23 residues displayed a capacity to solely form a 1:1 complex with NuMA, whereas the long-form AGS3-TPR also presented a weaker propensity compared to LGN to form high-order oligomeric complex with NuMA. Notably, Insc was capable of disrupting the oligomeric AGS3/NuMA complex, while the LGN/NuMA hexamer complex remained stable even in the presence of Insc (Fig. 7). These *in vitro* findings provide insights into the molecular mechanism underlying the distinct roles of LGN and AGS3 in regulating oriented cell divisions.

**Figure 7.**
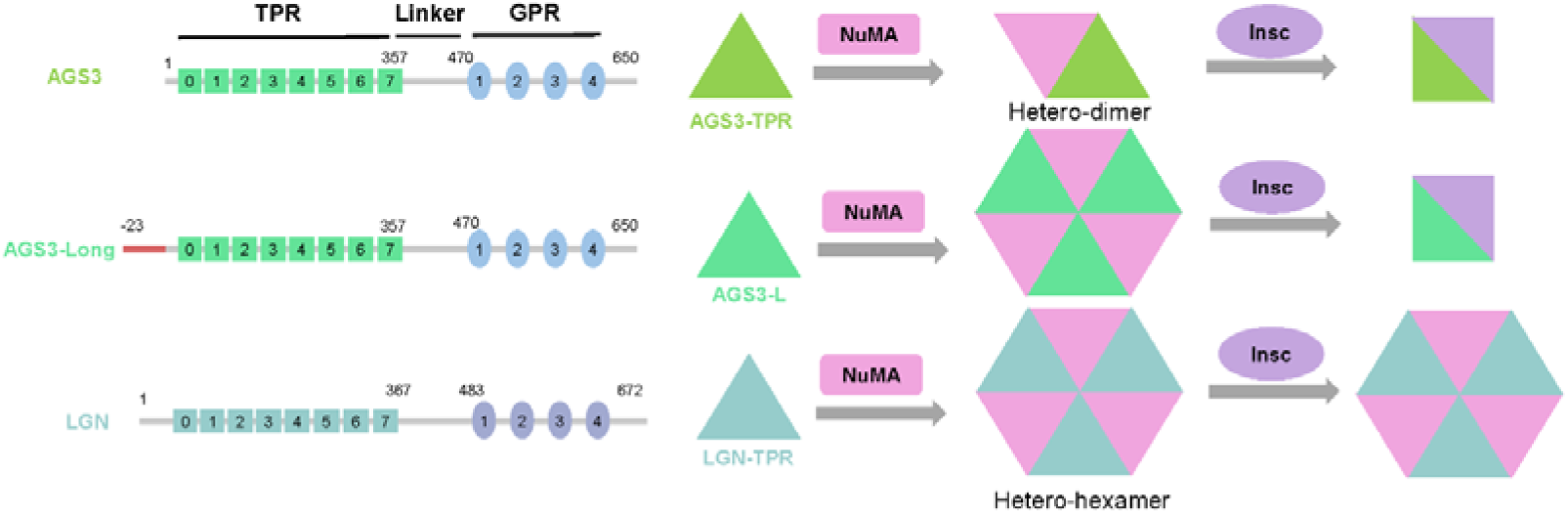
A model for the molecular mechanism underlying the distinct roles of LGN and AGS3 in regulating oriented cell divisions. The paralogs LGN and AGS3 share high sequence similarity and similar characteristics of partner interactions with NuMA and Insc. However, the multimeric AGS3/NuMA complex exhibits reduced stability compared to LGN/NuMA complex and is susceptible to disruption by Insc.

Our study corroborated previous observations that AGS3, due to its inability to form stable high-order oligomeric complexes with NuMA, cannot effectively regulate spindle orientation (Pirovano et al., 2019). This aligns with experimental evidence demonstrating that AGS3 could not rescue spindle orientation defects in LGN knockdown cells (Saadaoui et al., 2017). Our study demonstrated that AGS3-TPR as well as FL-AGS3 exhibit similar binding affinity to NuMA as those of LGN (Table 1 and Fig. S7) and can compete with LGN for NuMA binding (Fig. S9). Consequently, AGS3 most likely plays a role in reducing the quantity of oligomeric LGN/NuMA complex responsible for spindle orientation function, supporting the observation that AGS3 functions antagonistically to LGN in inhibiting the perpendicular divisions (Descovich et al., 2023).

It is noteworthy to emphasize that both full-length LGN and AGS3 exhibit a significantly reduced capacity to form oligomeric complexes with NuMA, even in the open conformation induced by the 4E mutation. The binding affinity to NuMA seems to be unaffected by Gα_i3_·GDP, suggesting that Gα_i3_·GDP is unlikely to facilitate the release of the auto-inhibited conformation. Another significant regulatory factor influencing the formation of oligomeric complexes between LGN or AGS3 and NuMA is Insc, which has the ability to disrupt the hetero-hexamer complex of AGS3-L/NuMA. Insc is known to localize to the apical cortex during asymmetric cell division(Kamakura et al., 2013). Although AGS3 has limited ability to form oligomeric complex with NuMA, Insc may also play an additional role in hindering the formation of a functional complex between AGS3 and NuMA for spindle orientation. Furthermore, Insc may exhibit a preference for binding to AGS3 over LGN, given that the oligomeric LGN/NuMA complex is resistant to Insc binding. As a result, AGS3 could potentially compete with LGN for Insc binding, and effectively sequestering LGN away from the apical cortex. This aligns with the observation that overexpression of AGS3 could weaken the polarized distribution of LGN (Descovich et al., 2023).

In the present study, we did not identify any functional disparities between the GoLoco domains of AGS3 and LGN *in vitro*. The individual GL motifs, the entire GoLoco domains and the full-length proteins of AGS3 and LGN all demonstrate analogous binding affinities to Gα_i3_·GDP. These results suggest that AGS3 and LGN likely possess comparable affinities towards Gαi *in vivo* as well. Consequently, the distinctive cellular localization patterns observed for AGS3 and LGN are unlikely to be primarily attributable to differences in their interactions with Gαi. It is conceivable that additional proteins may play a role in regulating the localization dynamics of LGN and AGS3 within cell. Further investigation into these additional regulatory factors could provide deeper insights into the mechanisms governing the spatial distribution and functional activities of LGN and AGS3 in cellular contexts.

## Materials and methods

### Protein expression and purification

The TPR domains of mouse LGN (1-409), mouse AGS3 (1-405 or -23-405), the GoLoco domains of LGN (483-650) and AGS3 (470-631), and the N-terminal fragment of mouse Insc (18-66), as well as the C-terminal fragments of human NuMA (1886-1912 or 1808-2001), were individually inserted into a modified version of the pET32a vector, pET28a-sumo or pGEX-6P-1 vector. Consequently, the resultant proteins featured a trx tag, sumo tag or GST tag at the N-termini. Human Gα_i3_ was cloned into a pET-M3C vector, resulting in a protein with a His6 tag at the N-termini. The FL-AGS3/LGN fusion proteins were cloned into a modified version of pET32a vector. All mutations were generated through standard PCR-based mutagenesis method and confirmed via DNA sequencing. Subsequently, the recombinant proteins were expressed in *E. coli* BL21 (DE3) host cells at 16 LJ and were purified using Ni^2+^-NTA or GST-agarose affinity chromatography followed by size-exclusion chromatography.

### GST pull-down assay

GST or GST-tagged proteins (final concentration: 4 μM) were initially applied to 50 μL GSH-Sepharose 4B slurry beads in a 500 μL assay buffer containing 50 mM Tris (pH 8.0), 100 mM NaCl, 1 mM 2-mercaptoethanol and 1 mM EDTA. Subsequent to four times of buffer washing, the beads loaded with GST fusion proteins were then mixed with potential binding partners (final concentration: 8 μM), and the mixtures were incubated for 1 hour at 4 LJ. After four additional washing steps, proteins captured by the affinity beads were eluted by boiling, separated via 12% SDS–PAGE, and visualized through Coomassie blue staining.

### Isothermal titration calorimetry

ITC measurements were performed using a Malvern PEAQ-ITC Micro calorimeter (Micro Cal) at 25 °C. All protein samples were dissolved in a buffer containing 50 mM Tris (pH 8.0), 100 mM NaCl, and 1 mM EDTA. For AGS3-TPR or AGS3-L protein, the salt concentration of NaCl is increased to 300 mM for optimal solubility. Titrations were carried out by injecting 40 μL aliquots of one protein (0.2 mM) into another protein (0.02 mM). The injections were spaced at 1 or 2-minute intervals to ensure that the titration peak returned to the baseline. The titration data were analyzed using the Origin software provided by Microcal and fitted with the one-site binding model.

### Analytical size-exclusion chromatography

An AKTA FPLC system (GE Healthcare) was employed to conduct analytical size exclusion chromatography (SEC). Protein samples were applied onto a Superose 12 10/300 GL column (GE Healthcare), which had been pre-equilibrated with a buffer containing 50 mM Tris-HCl (pH 8.0), 100 mM NaCl, 1 mM 2-mercaptoethanol, and 1 mM EDTA.

### Static Light Scattering

Protein characterization was accomplished through size exclusion chromatography coupled to multi-angle light scattering (SEC-MALS). Multi-angle light scattering was conducted using the HELEOSLJ instrument from Wyatt Technology. Generally, 100 μL of concentrated purified samples, with an approximate concentration of 10 μM, were applied to the columns.

## Acknowledgements

This work was supported by National Science Foundation of China (22073018, 22377015).

## Conflict of interest

The authors declare that they have no conflicts of interest with the contents of this article.

## Notes

### Competing Interest Statement

The authors have declared no competing interest.

