## Supplementary Information for "Molecular Insights into AGS3’s Role in Spindle Orientation: A Biochemical Perspective"

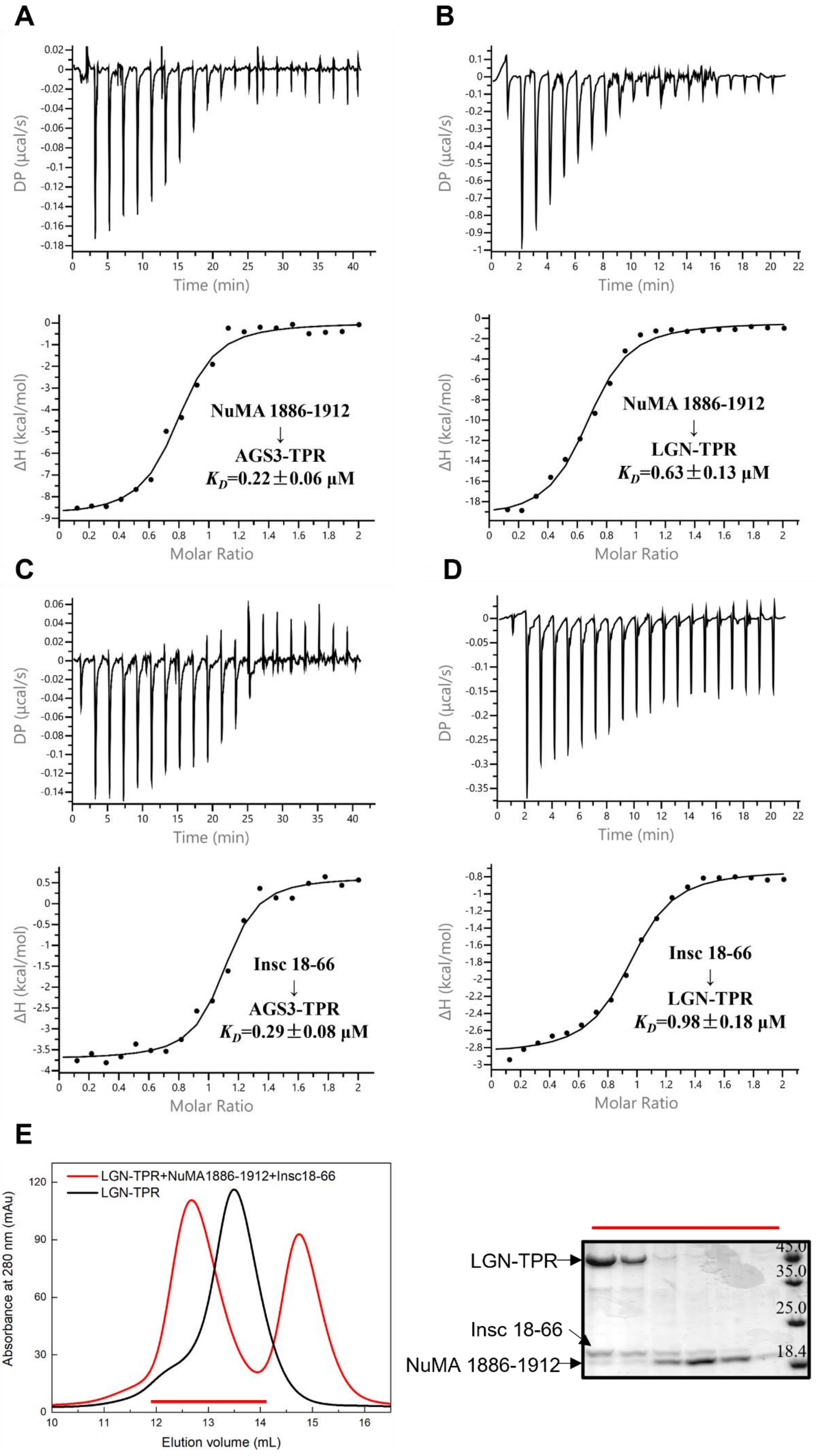
A
DP (µcal/s)
Time (min)
ΔH (kcal/mol)
Molar Ratio
NuMA 1886-1912
↓
AGS3-TPR
$K_D$=0.22±0.06 µM
B
DP (µcal/s)
Time (min)
ΔH (kcal/mol)
Molar Ratio
NuMA 1886-1912
↓
LGN-TPR
$K_D$=0.63±0.13 µM
C
DP (µcal/s)
Time (min)
ΔH (kcal/mol)
Molar Ratio
Insc 18-66
↓
AGS3-TPR
$K_D$=0.29±0.08 µM
D
DP (µcal/s)
Time (min)
ΔH (kcal/mol)
Molar Ratio
Insc 18-66
↓
LGN-TPR
$K_D$=0.98±0.18 µM
E
LGN-TPR+NuMA1886-1912+Insc18-66
LGN-TPR
Absorbance at 280 nm (mAu)
Elution volume (mL)
LGN-TPR
Insc 18-66
NuMA 1886-1912
45.0
35.0
25.0
18.4

**Figure S1.** AGS3-TPR binds to Insc and NuMA similarly to LGN-TPR. (A-D) ITC measurements of the bindings of AGS3-TPR/LGN-TPR protein to NuMA (1886-1912) and Insc (18-66). (E) The SEC analysis of the mixture of LGN-TPR, NuMA (1886-1912) and Insc (18-66) at a 1:2:2 ratio.

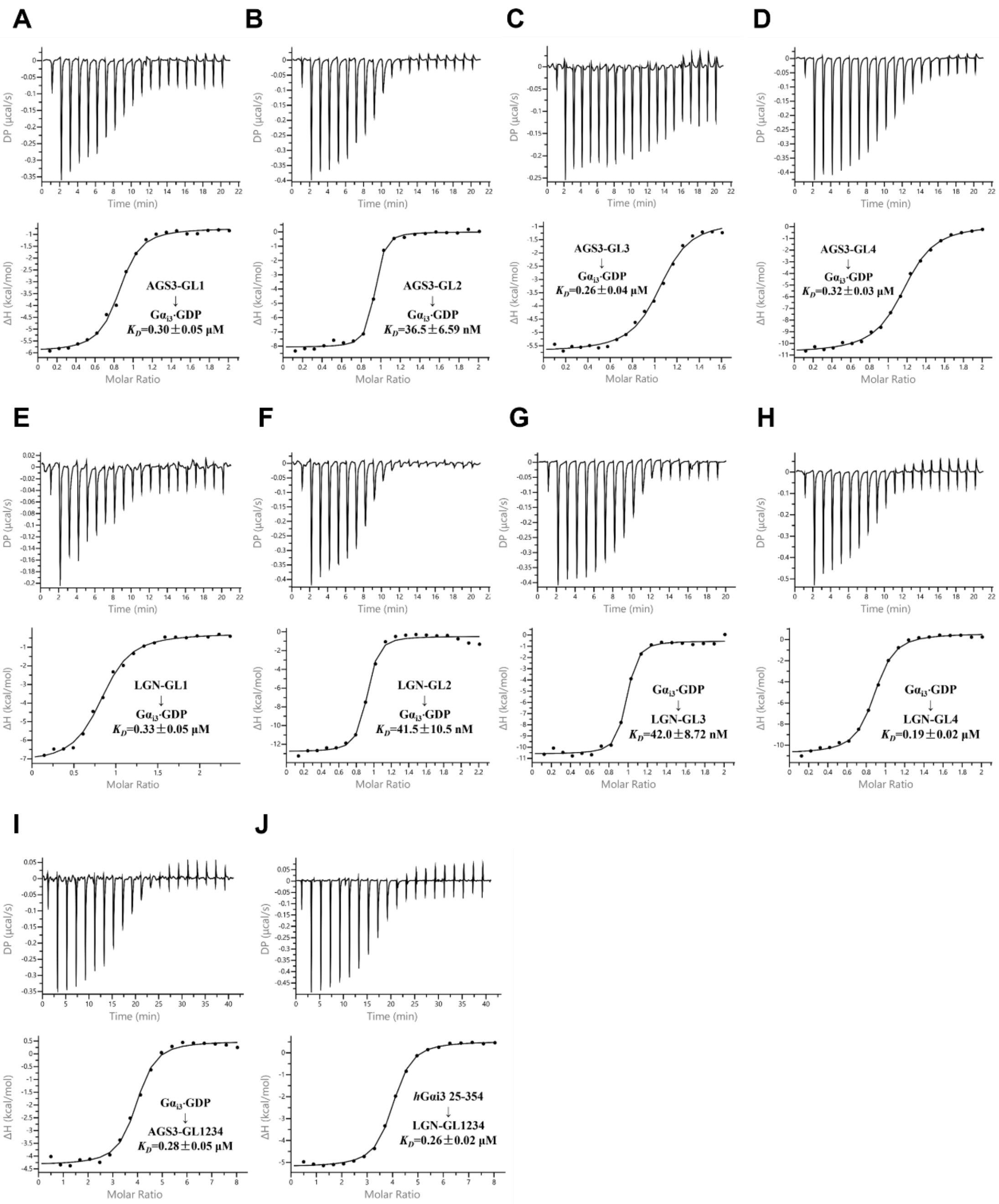


**Figure S2.** (A-D) ITC measurements of the bindings of $G\alpha_{i3}$·GDP to AGS3 GL motifs. (E-H) ITC measurements for the binding affinities of $G\alpha_{i3}$·GDP to LGN GL motifs. (I-J) ITC measurements for the binding affinities of $G\alpha_{i3}$·GDP to the GoLoco domains of AGS3 and LGN.

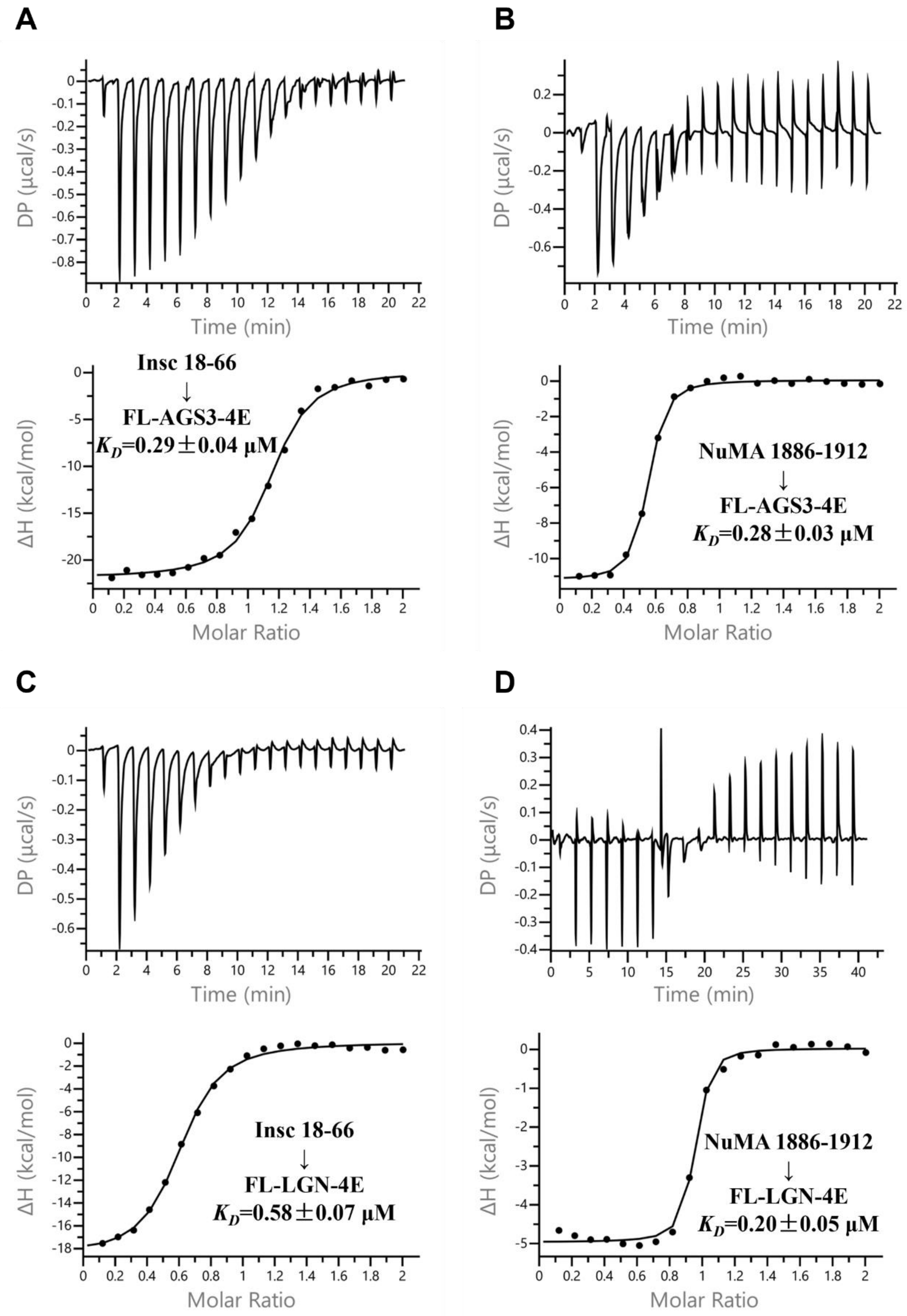


**Figure S3.** ITC measurements of the bindings of FL-AGS3-4E or FL-LGN-4E protein to Insc (18-66) and NuMA (1886-1912).

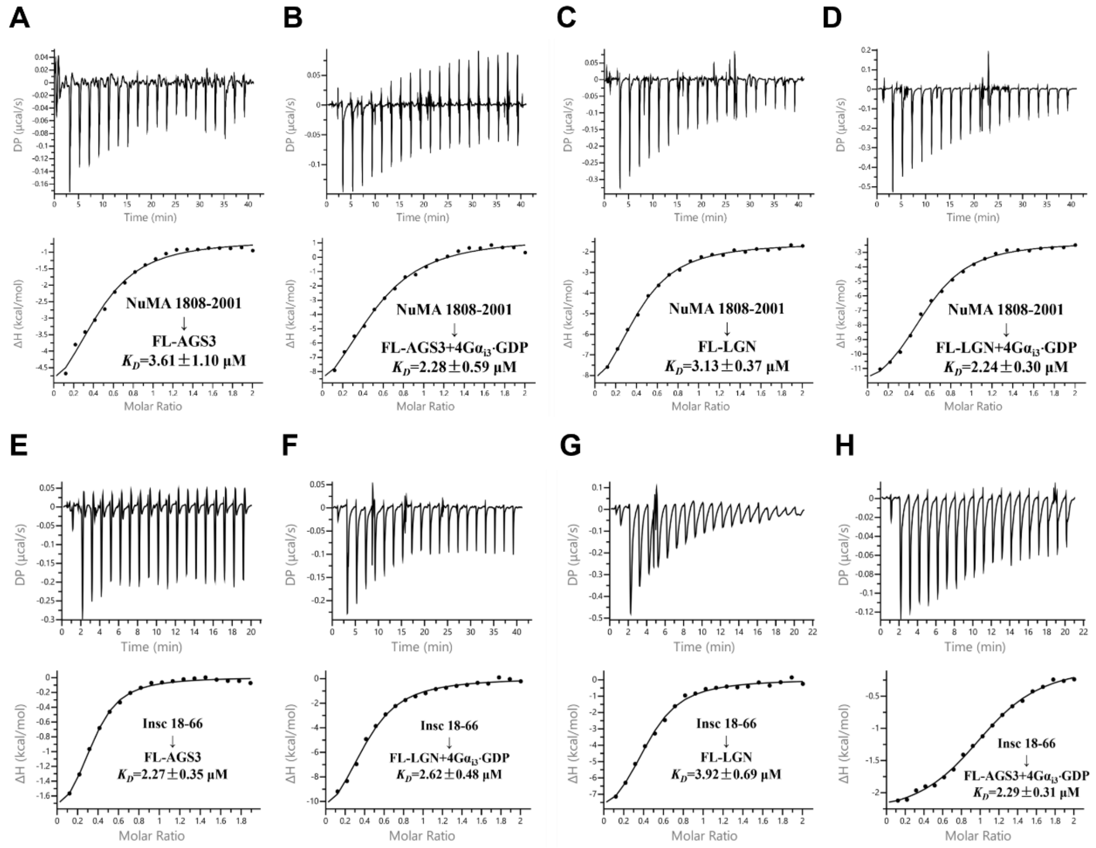


**Figure S4.** (A-D) The ITC measurements of the interactions between NuMA (1808-2001) and FL-AGS3/FL-LGN protein in the presence or absence of G$\alpha_{i3}$·GDP. (E-H) The ITC measurements of the interactions between Insc (18-66) and FL-AGS3/FL-LGN in the presence or absence of G$\alpha_{i3}$·GDP.

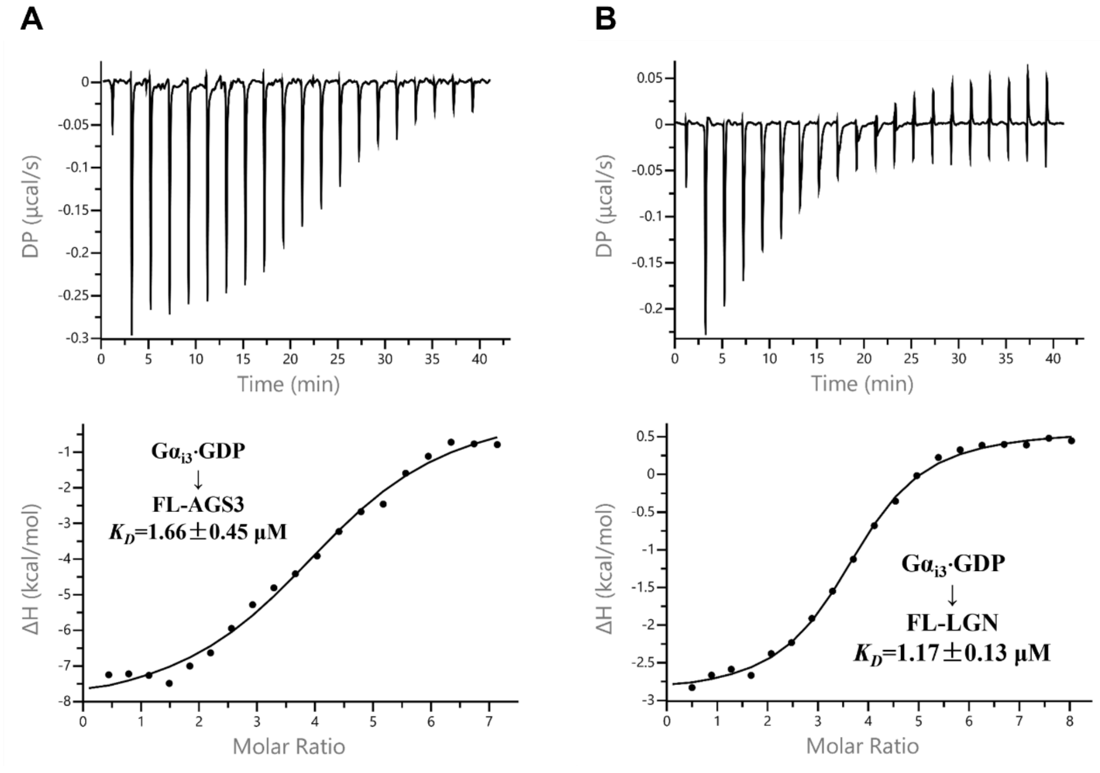


**Figure S5.** The ITC measurements for the interactions between $G\alpha_{i3}$·GDP and FL-AGS3 (A) or FL-LGN (B).

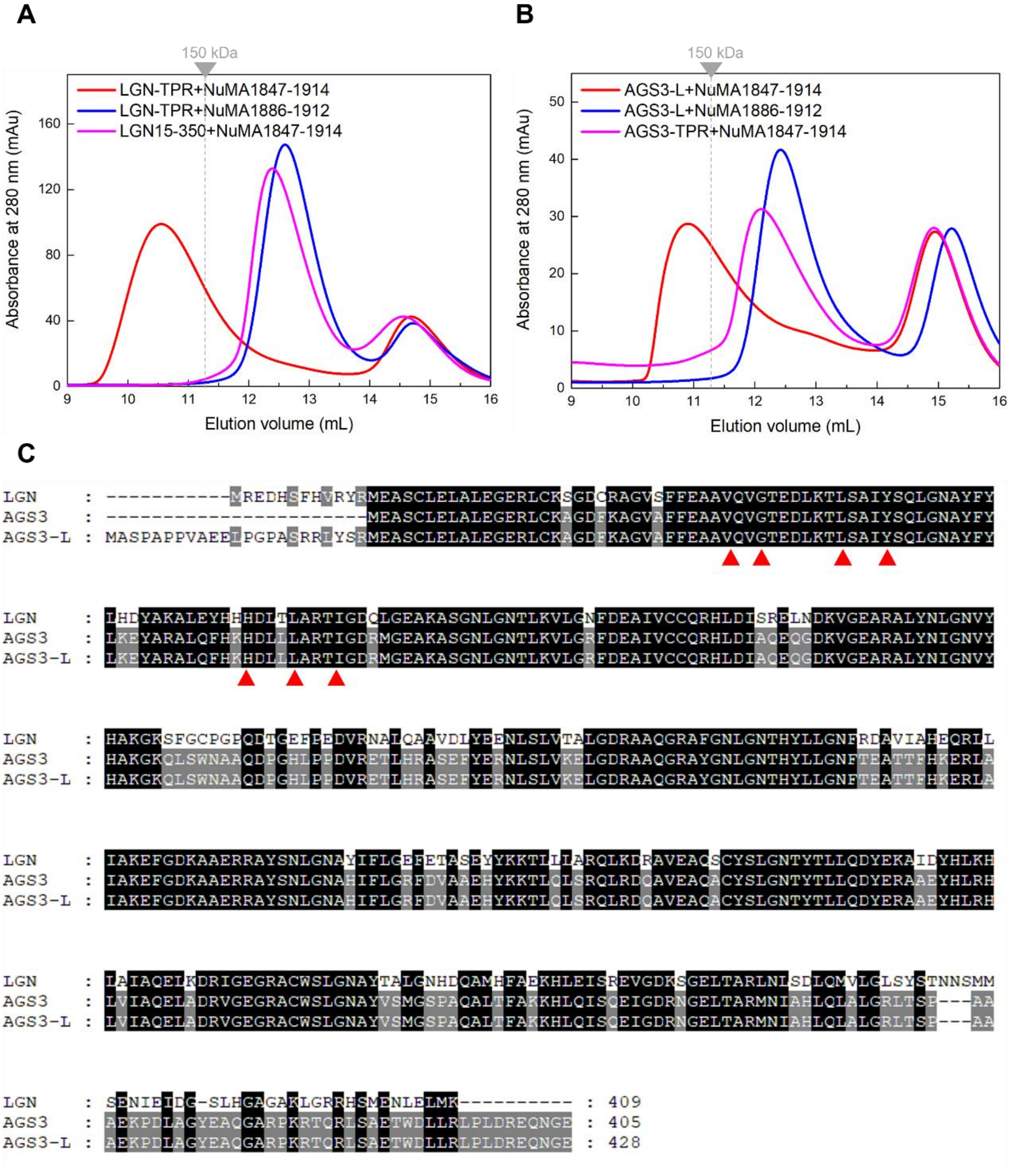


**Figure S6.** (A) SEC analyses of the complexes formed between various fragments of LGN-TPR and NuMA (1847-1914) or NuMA (1886-1912). (B) SEC analyses of the complexes formed between various fragments of AGS3-TPR and NuMA (1847-1914) or NuMA (1886-1912). (C) Sequence alignment of the N-terminal regions of LGN and AGS3 isoforms. The long isoform AGS3-L has additional 23 residues in the N-terminal which shows low homology comparing to LGN. The red triangles indicate the conserved residues in TPR that interact with NuMA (1847-1861).

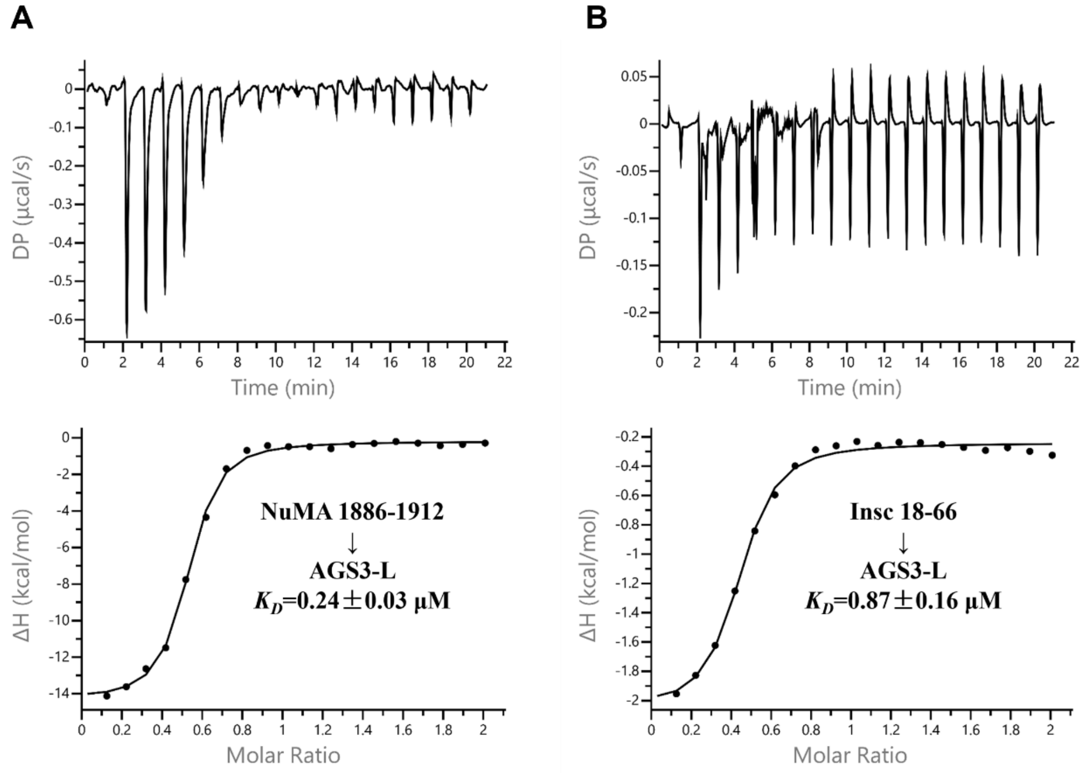


**Figure S7.** ITC measurements for the interactions between AGS3-L and NuMA (1886-1912) (A) or Insc (18-66) (B).

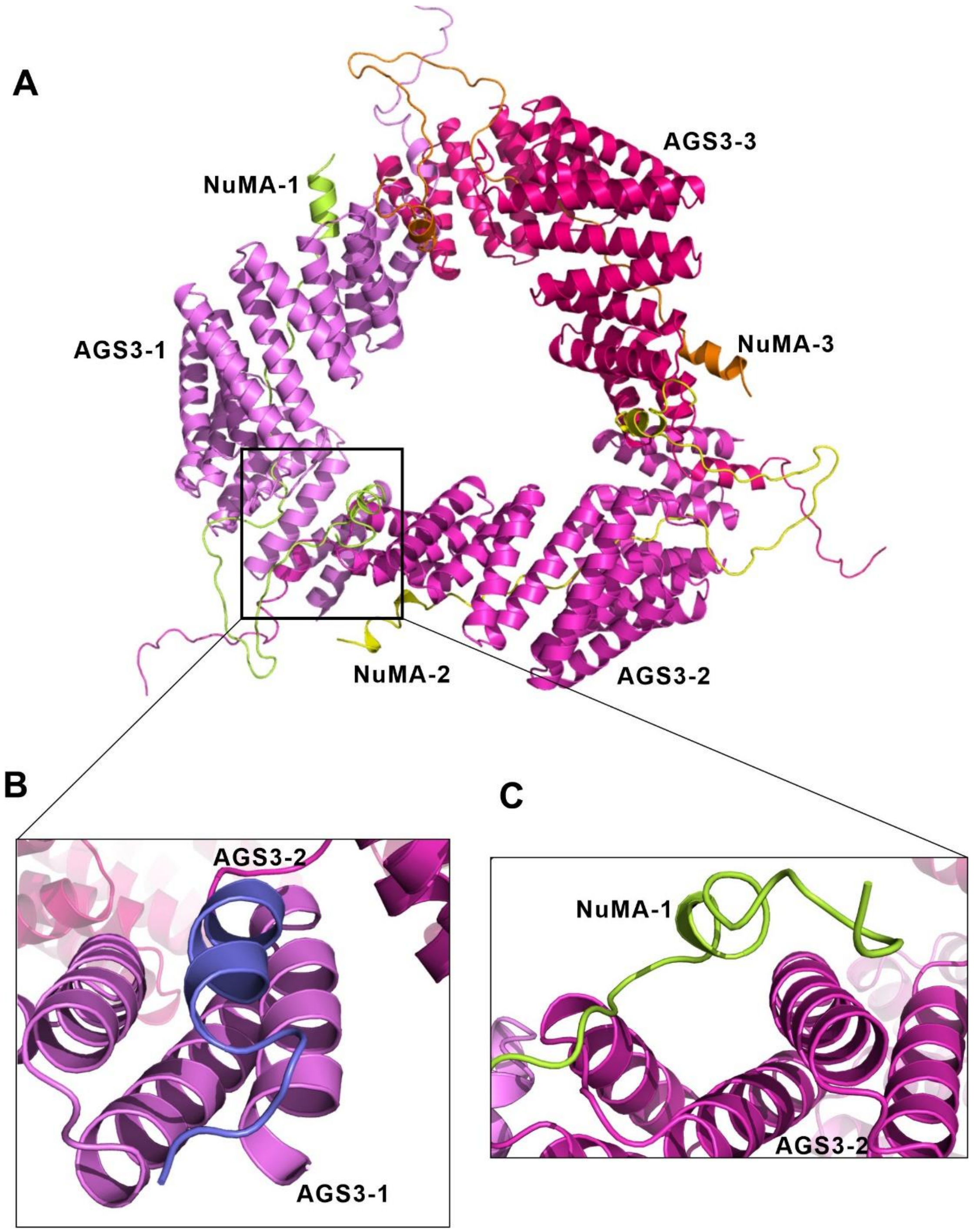


**Figure S8.** (A) The structural model of AGS3-L/NuMA hexamer complex. (B) The N-terminal of AGS3-L interacts with the adjacent AGS3 TPR8 to form a four-helix bundle. The -23 to -1 residues of AGS3-1 are highlighted in slate blue. (C) The N-terminal part of NuMA-1 interacts with the neighboring AGS3-2 to facilitate the hexamer formation.

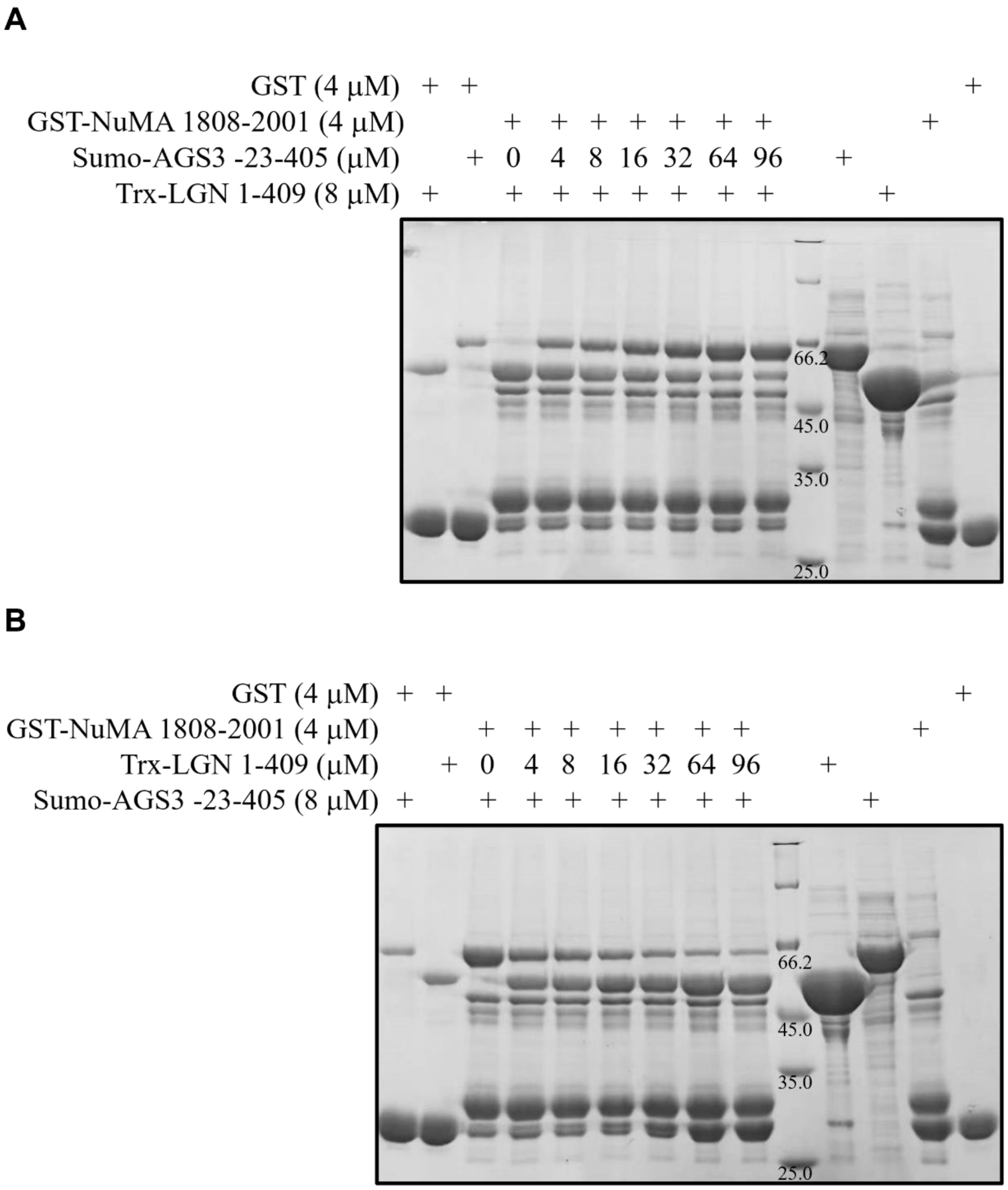


**Figure S9.** (A) GST Pull-down assay shows that with the increase of AGS3-L, the amount of NuMA-bound LGN-TPR decreased. (B) GST Pull-down assay shows that with the increase of LGN-TPR, the amount of NuMA-bound AGS3-L decreased.
